# Modelling the spatial dynamics of oncolytic virotherapy in the presence of virus-resistant tumor cells

**DOI:** 10.1101/2022.04.06.487254

**Authors:** Darshak K. Bhatt, Thijs Janzen, Toos Daemen, Franz J. Weissing

## Abstract

Oncolytic virotherapy is a promising form of cancer treatment that uses native or genetically engineered viruses to target, infect and kill cancer cells. Unfortunately, this form of therapy is not effective in a substantial proportion of cancer patients, partly due to the occurrence of infection-resistant tumor cells. To shed new light on the mechanisms underlying therapeutic failure and to discover strategies that improve therapeutic efficacy we designed a cell-based model of viral infection. The model allows to investigate the dynamics of infection-sensitive and infection-resistant cells in tumor tissue in presence of the virus. To reflect the importance of the spatial configuration of the tumor on the efficacy of virotherapy, we compare three variants of the model: two 2D models of a monolayer of tumor cells and a 3D model. In all model variants, we systematically investigate how the therapeutic outcome is affected by properties of the virus (e.g. the rate of viral spread), the tumor (e.g. production rate of resistant cells, cost of resistance), the healthy stromal cells (e.g. degree of resistance to virus) and the timing of treatment. We find that various therapeutic outcomes are possible when resistant cancer cells arise at low frequency in the tumor. These outcomes depend in an intricate but predictable way on the death rate of infected cells, where faster death leads to rapid virus clearance and cancer persistence. Our simulations reveal three different causes of therapy failure: rapid clearance of the virus, rapid selection of resistant cancer cells, and a low rate of viral spread due to the presence of infection-resistant healthy cells. Our models suggest that improved therapeutic efficacy can be achieved by sensitizing healthy stromal cells to infection, although this remedy has to be weighed against the toxicity induced in the healthy tissue.

## Introduction

Advances in genetic engineering and synthetic biology have allowed the rational design of safer and more efficient anti-cancer biotherapies [1,2]. Oncolytic viruses belong to such a class of therapeutics, where native or genetically modified viruses are employed as agents to preferentially target, infect and kill cancer cells [2,3]. This strategy is based on the rationale that cancer cells often harbor mutations in innate antiviral responses, which makes them a sensitive target of oncolytic virotherapy [4,5]. To this end, a wide range of viral vectors have been tested in clinical trials suggesting promising therapeutic benefit [6]. Unfortunately, only a small proportion of patients treated with oncolytic viruses were found to have a long-term benefit [7]. Potential explanations could be the technical limitations related to virus delivery and treatment, variability in cancer types, generation of weak anti-tumor immunity, individual differences among patients, and occurrence of infection-resistant cancer cells [4,8–10]. Here we focus on the role of resistance, as it remains to be a relatively unexplored area of research[11].

The presence of resistant cancer cells in the tumor tissue may restrict the efficient spread of the virus and thereby undermine its oncolytic potential. The therapeutic efficacy of a virus does therefore not only depend on the properties of the virus, but also on the density and spatial configuration of resistant tumor cells, as these factors are crucial for the spatial dynamics of viral spread in the tumor. To unravel the causes of therapeutic resistance towards virotherapy it is therefore paramount to map the spatial interactions between virus, infection-sensitive cancer cells and virus-resistant cancer cells, and the implications of these interactions for tumor eradication. This can be done with experimental approaches using in-vitro assays or animal models to investigate the complex spatial dynamics that occur between virus and the different cell types present in the tumor. However, technological limitations do not yet allow the large-scale screening that would be required to get a full picture of when, how and why the complex interplay of multiple factors results in therapeutic failure. In a situation like this, mathematical or computational models can be useful by providing insights into the role and relative importance of the factors governing the spatial dynamics of virotherapy in a tumor, and by suggesting adaptations of the therapy that can subsequently be scrutinized experimentally.

Various mathematical and computational models have been developed to elucidate the outcome of virus-tumor interactions, and some have even explicitly considered the geometry of cells and the spatial nature of the tumor tissue in the model design [12–14]. For example, Berg and colleagues [15] developed two- (2D) and three-dimensional (3D) spatial models that are informed by in vitro experimental data of virus infection in a monolayer of cells (2D) and tumor spheroids (3D), respectively. They demonstrated that the introduction of a third dimension alters the dynamics of virus-tumor interaction and significantly influences the ability of the virus to eradicate the tumor. Another study considering the spatial dynamics of virus-tumor interactions [16] demonstrated that there is a dichotomy in therapeutic outcomes due to antiviral signaling mediated by induction of interferon production in infected cells. So far, this has been the only modelling study that addresses the possible role of resistance to viral infection. However, to our knowledge, no study has included resistant cancer cells explicitly in a spatial model, thus allowing one to assess the impact of the presence and spatial configuration of such cells on the viral dynamics and efficacy of virotherapy.

We therefore set out to develop such a model. The model considers a tumor tissue with different cell types that proliferate and die with different rates and that are susceptible to viral infection. As it is known that the spatial configuration of the tumor tissue can play a key role in determining the therapeutic efficacy of oncolytic viruses [15], we systematically compare the outcome of three model variants, which consider a regular 2D monolayer, a more natural Voronoi 2D monolayer, and a 3D configuration of tumor cells. In each of these model variants, we aim to understand the interplay of viral and tumor dynamics and, in particular, the factors determining the persistence of resistant cancer cells that underlies therapeutic resistance.

We also study the role of healthy stromal cells (such as cancer-associated fibroblasts, epithelial cells and endothelial cells) regarding the efficacy of virotherapy as these cells have functional innate immune responses and are resistant to viral infection [9,10,17]. To limit the complexity of our models, we do not consider stromal cell mediated immune responses, such as antiviral signaling and myeloid or lymphoid anti-tumor responses. In the present study, we focus on the role of stromal cells regarding spread of the virus, and we investigate whether and when sensitizing healthy stromal cells can enhance persistence of the virus and thus improve the therapeutic outcome.

Thus, we systematically assess the impact of virus spread and oncolysis, resistance mediated by cancer and stromal cells, frequency of resistant cancer cells and cost of resistance on therapeutic outcomes. Furthermore, we consider sensitization of stromal cells towards viral infection and the possibility of virus dispersal in the tumor as possible strategies to improve virotherapy. Finally, by comparing the 2D and 3D models we aim to gain an insight on the spatial dynamics of virotherapy so that it can be rationally optimized for better outcomes.

## Model description

### Overview of the model

Our model takes the work of Berg *et al*. [15] as point of departure. We model the growth of tumor and stromal cells on a cell-based spatial grid, using an event-based time structure. Whereas the grid remains static, cell types can proliferate across the grid, reflecting growth of both stromal and cancer cells. Cancer cells can become infected by an oncolytic virus, which is programmed to preferentially target and kill cancer cells while sparing stromal cells. Upon proliferation, tumor cells are able to acquire resistance against the oncolytic virus, which may come at a cost and may alter the reproductive potential of a cell to have a slower proliferation rate or a faster death rate. We explore how this cost of resistance influences the efficacy of the virus. Furthermore, we explore how the sensitivity of stromal cells to accidental infection by the oncolytic virus influences therapeutic outcomes.

### Spatial organization of the tumor

The model consists of a 2-dimensional or a 3-dimensional grid, consisting of nodes (e.g. the cells) connected with their neighbors, these connections reflect the cell’s boundaries. Cells at the edges of the grid have fewer neighbors than cells in the interior. In two dimensions, we consider either a regular lattice (figure 1A), where cells interact with their four direct neighbors (horizontally and vertically), or a lattice resulting from a Voronoi tessellation (figure 1B). The Voronoi tessellation is obtained by populating the spatial area with *N* node centers (where *N* denotes the total number of cells in the simulation) that are randomly distributed over the whole spatial area (their coordinates are drawn from a uniform distribution). Using Fortune’s algorithm [18], we calculate the Voronoi diagram that specifies *N* regions (“Voronoi cells”) around the node centers, which consist of all those points that are closer to their node center than to any other node center. We then used these Voronoi cells to determine which nodes neighbor each other. For the three dimensional grid, we only considered a regular lattice (figure 1C), where cells interact with their six direct neighbors (two in each dimension). Extension towards a three dimensional Voronoi tessellation is straightforward but was not further explored, as it is computationally demanding.

**Figure 1:**
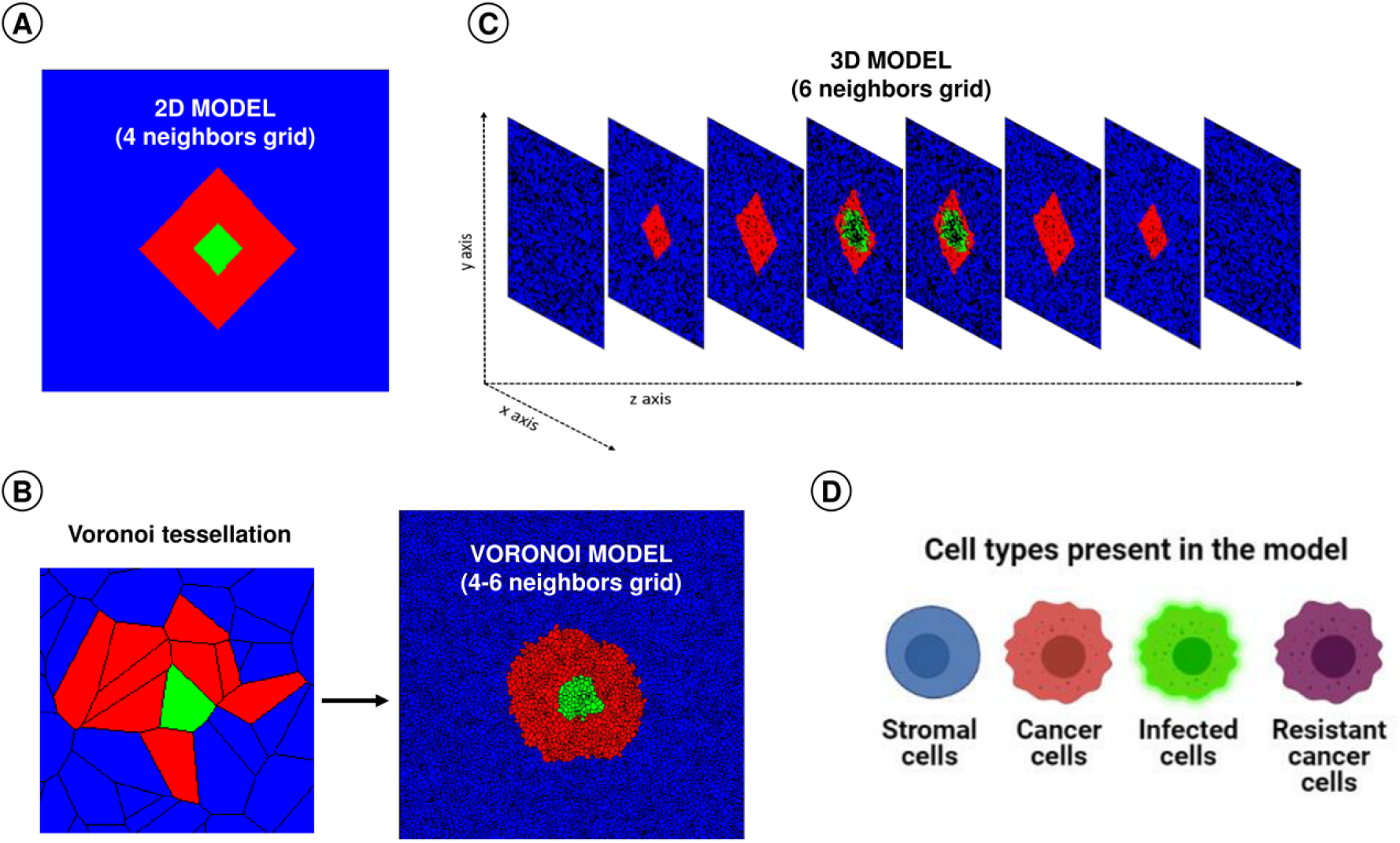
Visualization of the cell-based model variants. Spatial organization of (A) the 2-dimensional regular grid model, where each cell has 4 neighbors; (B) the 2-dimensional Voronoi grid model, where each cell has typically 4-6 neighbors; and (C) the 3-dimensional regular grid model, where each cell has 6 neighbors. (D) The four cell types in the model: stromal cells (blue), infection-sensitive cancer cells (red), virus-infected cancer cells (green) and resistant cancer cells (purple).

The grid thus generated was kept static throughout the simulation and only the identity of the nodes was allowed to change (see below for the associated cell types). Throughout the manuscript, we used a grid of 10,000 cells, such that in two dimensions, the grid was 100×100 cells (regular and Voronoi grid), or in three dimensions the grid was 22×22×22 cells. Supplementary figure 1 shows that the simulation outcome is only marginally affected by grid size, confirming that our choice of grid size provides representative results.

### Cell types present in the model

Each node in the grid can be inhabited by one of four different cell types: healthy (stromal) cells, infected (cancer or stromal) cells, and two different cancer cell types: infection-sensitive cancer cells and cancer cells resistant to virus infection (figure 1D). Stromal cells here represent cancer-associated fibroblasts, endothelial cells and epithelial cells, but do not include immune cells. Each node is either empty or occupied by one of the four cell types.

Stromal or cancer cells can proliferate (with rates *b*_*s*_ and *b*_*c*_ respectively) and divide into neighboring empty nodes. When multiple neighboring empty nodes are available, one is chosen at random. Virus-infected cells do not divide into empty nodes, but they can infect neighboring nodes inhabited by susceptible cancer cells (with rate *b*_*i*_). We also consider the possibility that stromal cells are infected by the virus. To this end, we assume that the susceptibility of a stromal cells is *S*_s_ times as large as the susceptibility of a cancer cell. A virus-infected cell infects a neighboring node with a probability that is identical with the susceptibility of this node, where the susceptibility of a susceptible cancer cell is given by 1 and the susceptibility of a stromal cell is given by *S*_s_. Each cell also has a fixed probability, independent of its neighbors, to die and leave an empty space in the grid (with rates *d*_*s*_, *d*_*c*_ and *d*_*i*_ for stromal, cancer and virus-infected cells respectively; in line with Berg *et al*. [15] we assume that virus-infected cancer cells and virus-infected stromal cells have the same death rate *d*_*i*_). The rates of proliferation and death of each cell type determine the probability of a cell to occupy a neighboring node. Cancer cells have a higher rate of proliferation and a lower death rate as compared to normal cells, defining their tumorigenic features. Upon proliferation into an empty cell, the resulting daughter cell of a virus-susceptible cancer cell can acquire resistance to the virus infection with a fixed probability *C*_*r*_. There may be a cost to this resistance, which is reflected by a different rate of proliferation and death for resistant cancer cells (*b*_*r*_ and *d*_*r*_ respectively).

The model was implemented as an event-based model, using a Gillespie algorithm [19]. All rates provided are in events per day, such that a rate of 2.0 reflects an event occurring twice per day, with an average waiting time of 0.5 days between events. Simulations were run until T_max_ days had passed, which we chose to be 1,000 days, as at this time point transient patterns had disappeared and firm conclusions could be drawn on the therapeutic outcome (figure 2).

**Figure 2:**
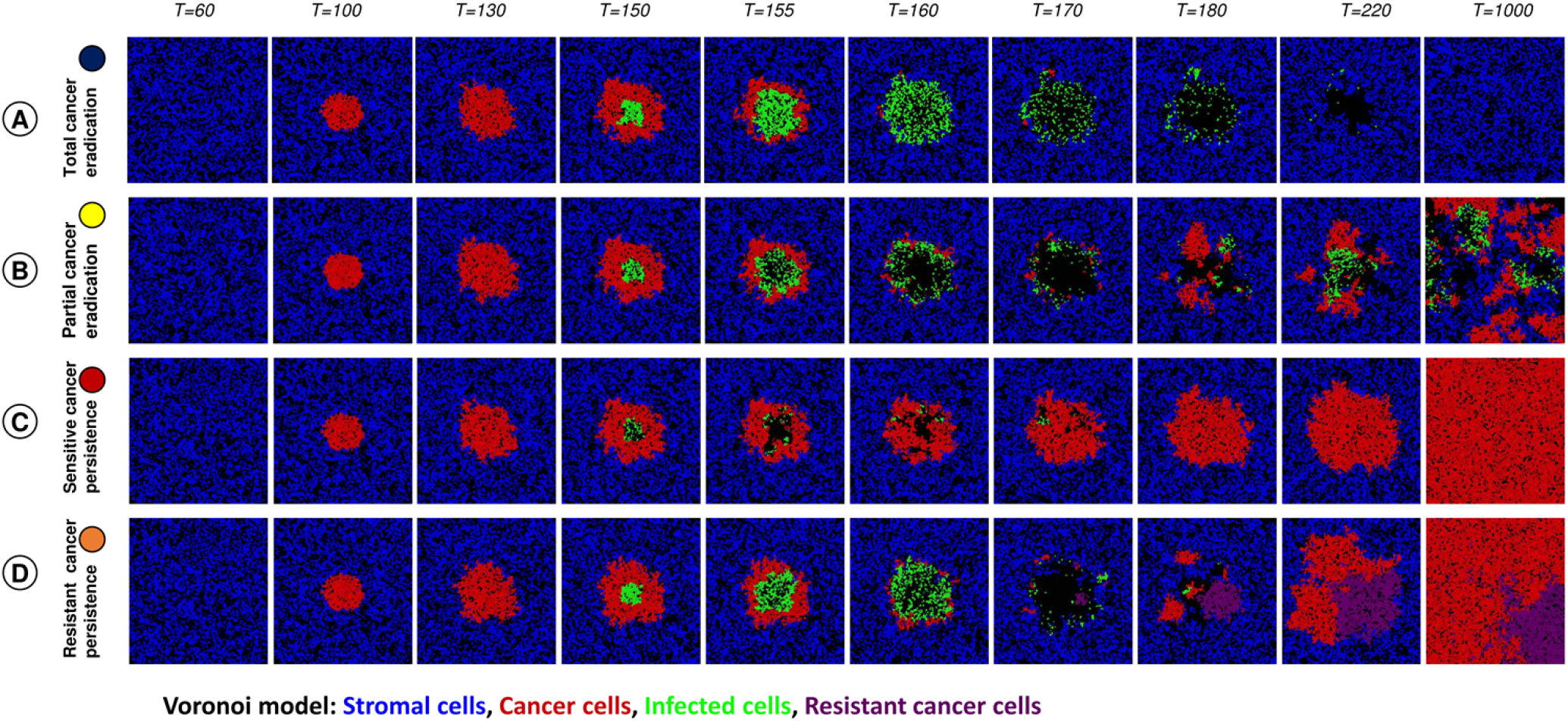
Visualization of different outcomes of oncolytic virotherapy. In our model, virus infection in the tumor tissue can result in four therapeutic outcomes: (A) total cancer eradication; (B) partial cancer eradication; (C) persistence of a virus-sensitive cancer; and (D) persistence of a partially virus-resistant cancer.

### Infection via virus diffusion

Apart from contact-based spread of virus from an infected cell to a neighboring susceptible cancer cell (or a stromal cell), the model also considers the possibility of virus diffusion and spread after the death of an infected cell [20,21]. A dying cell might emit additional viral load upon lysis. In the model, a dying cell infects neighboring cells with a predefined probability (*P*_*id*_), up until the maximum distance (*D*_*i*_), where this distance is measured as the number of edges connecting the focal cells that face a probability of infection, and the dying cell. Thus, out of all susceptible neighbors within the maximum distance *D*_*i*_, a fraction *P*_*id*_ will become infected. We consider the direct neighbors of a dying cell to be at distance 1, the neighbors of the neighbors of this dying cell to be at distance 2, the neighbors of the neighbors of the neighbors of this dying cell to be at distance 3, and so on (see fig. 6B for an illustration). The infection of neighboring cells occurs immediately upon death and we do not explicitly model the diffusion of virus particles in order to retain simplicity of the model and computational tractability.

### Setup of a simulation

Initially stromal cells are introduced in the center of the simulation and allowed to fill the grid until an equilibrium is reached, such that the total number of stromal cells is dynamically stable in time, e.g. the entire grid is populated with the maximum density of normal cells. Cancer cells are introduced in the center of the grid and allowed to proliferate until time T_i_, after which the virus is introduced in the center of the tumor. For all simulations, virotherapy is only carried out once, where a fraction *F*_*i*_ of the cancer cells are infected. To do so, the central tumor cell is determined, and is infected. Then, all neighboring cells are infected until a fraction *F*_*i*_ of the cancer cells are infected. The introduction of the virus in the center of the cancer cells reflects virus injection within the tumor body. We also considered other infection routines (infection in the periphery, infection at random), but as they all gave similar outcomes (see Supplementary figure 2 and also observed by Wein et al. [22]), no results on these routines will be reported in this study.

## Results

As various parameters pertaining to the virus, cancer cells and stromal cells influence the dynamics of virus-tumor interaction, we systematically study their influence on therapeutic efficacy of oncolytic viruses. To this end, we first categorize the result of each simulation into a therapeutic outcome based on the cell types present at the end of the simulation.

### Classification of therapeutic outcomes

Figure 2 visualizes that a simulation can result in four therapeutic outcomes. (A) <u>Total cancer eradication</u> occurs when virus therapy results in the extinction of all tumor cells. In this case, the virus infection spreads rapidly through the tumor. After the virus-infected cells have died, only stromal cells persist in the end (figure 2A). (B) <u>Partial cancer eradication</u> occurs when stromal cells, cancer cells and infected cells coexist for a long period of time in a dynamically changing spatial configuration (figure 2B). In some cases, stromal cells go extinct due to their slower growth rate and only infected and uninfected cancer cells coexist. As infected cancer cells are “handicapped” and do not contribute to tumor formation, therapy is at least partly successful. (C) <u>Persistence of a virus-susceptible cancer</u> occurs when the population of virus-infected cancer cells goes extinct. In the end, the tumor cells take over the population as they outgrow the stromal cells due to their faster growth rate (figure 2C). This outcome corresponds to therapeutic failure, but as the cancer cells remain virus-sensitive, a new round of treatment might be successful. (D) <u>Persistence of a partially virus-resistant cancer</u> occurs when the virus infection leads to the spread of infection-resistant cancer cells, which in turn results in the extinction of the virus. The resulting population typically consists of a mixture of virus-resistant and virus-sensitive cells (figure 2D). On a longer-term perspective, the sensitive cells will outcompete the resistant cells (because of their higher growth rate and/or lower death rate), resulting in scenario (C). Still, scenario (D) may be viewed as a more definite failure of therapy, as a new round of treatment will have a low probability of success as long as a substantial fraction of the tumor consists of resistant cells. (E) Finally, there is also the possibility (not shown in figure 2) that the simulation results in the extinction of all cell types, although we do not consider this as a “therapeutic outcome” of our model.

### Outcome of virotherapy in the absence of resistant cancer cells

Figure 3A illustrates how, in the absence of resistant cancer cells, the therapeutic outcome is determined by the rate of viral spread (*b*_i_) and the death rate of infected cells (*d*_i_). The outcome largely depends on the ratio *d*_i_/*b*_i_ of these rates. Total cancer eradication (figure 3A, blue area) only occurs if the death rate of infected cells is an order of magnitude smaller than the rate of viral spread (small *d*_i_/*b*_i_). In this case, the virus can spread and eliminate all cancer cells. In contrast, a large ratio *d*_i_/*b*_i_ results in rapid viral clearance, persistence of the virus-susceptible cancer (figure 3A, red area) and hence in therapeutic failure. An intermediate ratio *d*_i_/*b*_i_ results in partial cancer eradication (figure 3A, yellow area). Interestingly, there is a small red “wedge” between the yellow and the blue area, indicating that total cancer eradication can also occur for relatively small values of *d*_i_/*b*_i_. We will discuss this phenomenon below.

**Figure 3:**
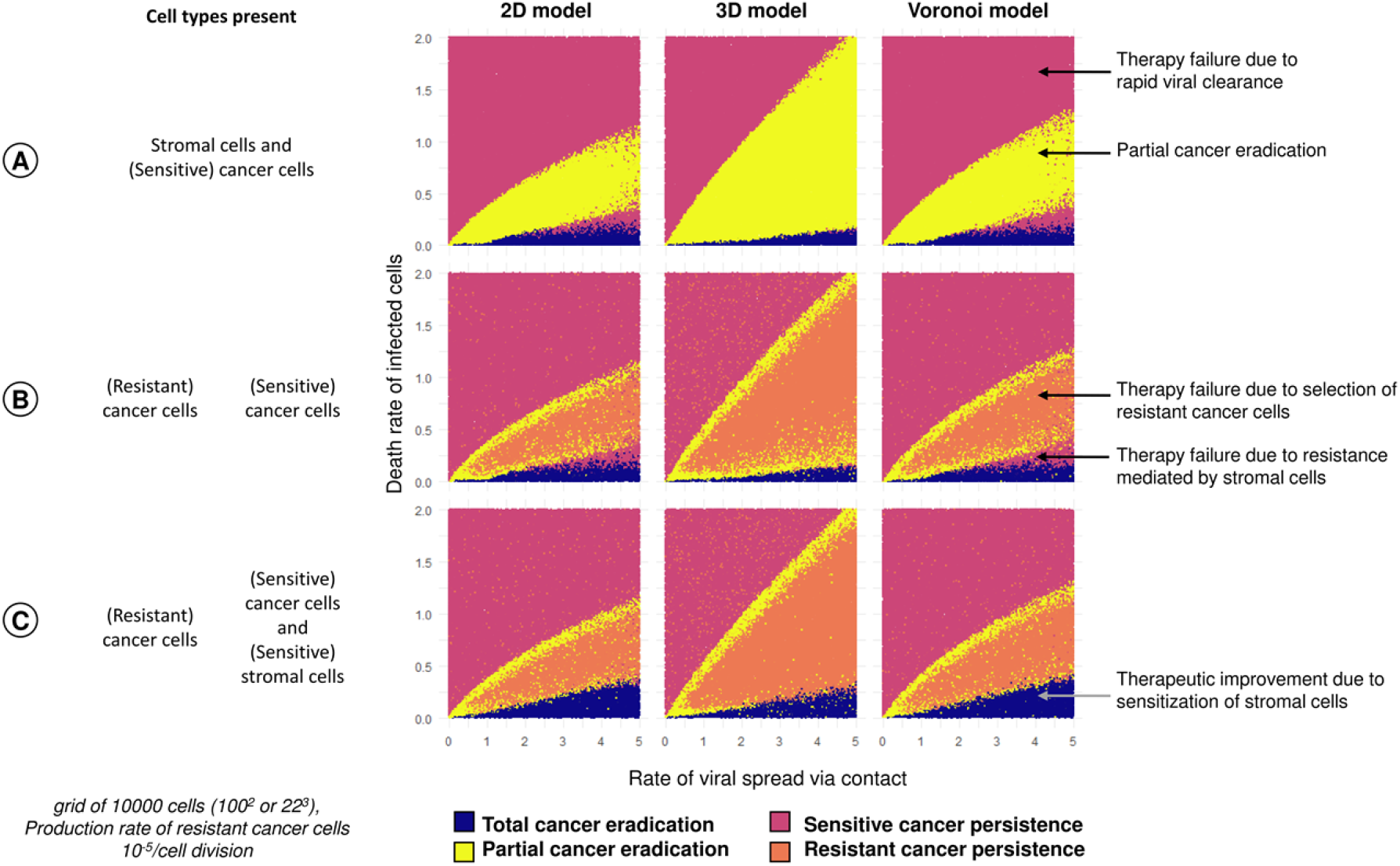
Effect of the rate of viral spread and death rate of infected cells on the outcome of virotherapy. For three scenarios: (A) absence of virus-resistant cancer cells; (B) production of resistant cancer cells with probability 10^−5^ per cell division; (C) presence of virus-sensitive stromal cells the panels show for three spatial configurations (regular 2D grid; regular 3D grid; 2D Voronoi tessellation) how the therapeutic outcome depends on the rates of viral spread (*b*_i_) and death rate of infected cells (*d*_i_). Each panel in the figure represents 100,000 simulations: for 100,000 parameter combinations (*b*_i_,*d*_i_) each simulation outcome is represented by a point, the color of which indicates the therapeutic outcome. Due to stochasticity, neighboring parameter combinations can differ in therapeutic outcome.

Qualitatively, the dependence of the therapeutic outcome on the ratio of the death rate of infected cells and the rate of viral spread is the same in all three spatial variants of our model. However, in the 3D model partial cancer eradication is observed in a broader parameter range than in the two 2D models (regular 2D and Voronoi). Moreover, the red wedge is smaller in the 3D model. These results are in good agreement with the conclusions drawn from the model developed by Berg et al [15]. Moreover, it is reassuring that our model reproduces crucial aspects of the virus-cancer dynamics (ring-spread or disperse-spread) that was experimentally observed and reproduced by a discrete-time agent-based model by Wodarz et al [14] (supplementary figure 3).

### Influence of resistant cancer cells on therapeutic outcomes

The inclusion of virus-resistant cancer cells is a novel aspect of our model. Figure 3B illustrates the effect of such cells on the therapeutic outcome when resistant cancer cells arise (by mutation) with a frequency of 10^−5^ per cancer cell division. In the presence of resistant cells, partial cancer eradication (yellow area) is no longer a prominent outcome. Instead, intermediate ratios *d*_i_/*b*_i_ more typically result in the persistence of a partially resistant cancer (orange area) and, hence, in total therapy failure.

Figure 3 shows the therapeutic outcome for a relatively small range of virus-induced spread and death rates of infected cells. Supplementary figure 4 presents the outcome of simulations for a much larger range of viral spread rates (0 to 25) and death rate of infected cells (0 to 5). This figure shows that for large infection rates (*b*_i_ > 10), partial tumor eradication (yellow area) and the persistence of partially resistant cancer (orange area) do no longer occur. Accordingly, the occurrence of resistant cancer cells is of minor therapeutic relevance if the viral spread rate is very large.

### Influence of stromal cells on therapeutic outcomes

In the default version of our model, we assume that virotherapy is designed to specifically infect and kill cancer cells, while healthy stromal cells are immune against viral infection. As a consequence, stromal cells may act as a spatial barrier, preventing the efficient spread of the virus in the tumor. We hypothesized that this barrier effect is responsible for therapeutic failure in the “red wedge” set of parameters in figures 3A and 3B (the red area between the blue area corresponding to total cancer eradication and the yellow/orange area corresponding to the partial eradication of virus-sensitive cancer cells). To test this hypothesis, we “sensitized” stromal cells, by allowing them to be infected at a similar rate as sensitive cancer cells. Figure 3C shows that, in line with our hypothesis, the “red wedge” is indeed removed by this intervention. In other words, virotherapy with a virus that can infect stromal cells as well as cancer cells can result in total cancer eradication in a situation where the cancer would persist if the case of virus-resistant stromal cells. However, the parameter range where the virus sensitivity of stromal cells is beneficial for the therapeutic outcome is relatively small. Moreover, the virus sensitivity of stromal cells comes at a cost, also in situations where it does not improve the therapeutic outcome. This is shown in supplementary figure 5b: in all scenarios, a smaller population of stromal cells remains when these cells are virus-sensitive. Additionally, when resistant cancer cells can persist, their number is typically positively affected by the presence of virus-sensitive stromal cells, as the presence of these cells increases the selective advantage of resistance (supplementary figure 5c).

### Factors influencing persistence of resistant cancer cells

Figure 4A shows how the likelihood of the four therapeutic outcomes is affected by the probability *C*_r_ that a virus-sensitive cancer cell acquires resistance due to mutation (see supplementary fig. 6 for a more detailed account). When the production rate of resistant cancer cells is smaller than 10^−5^ (the situation considered in Fig. 3B), virotherapy only rarely results in the persistence of (partially) resistant cancer; while this is the typical outcome when *C*_r_ is 10^−3^ or larger.

**Figure 4:**
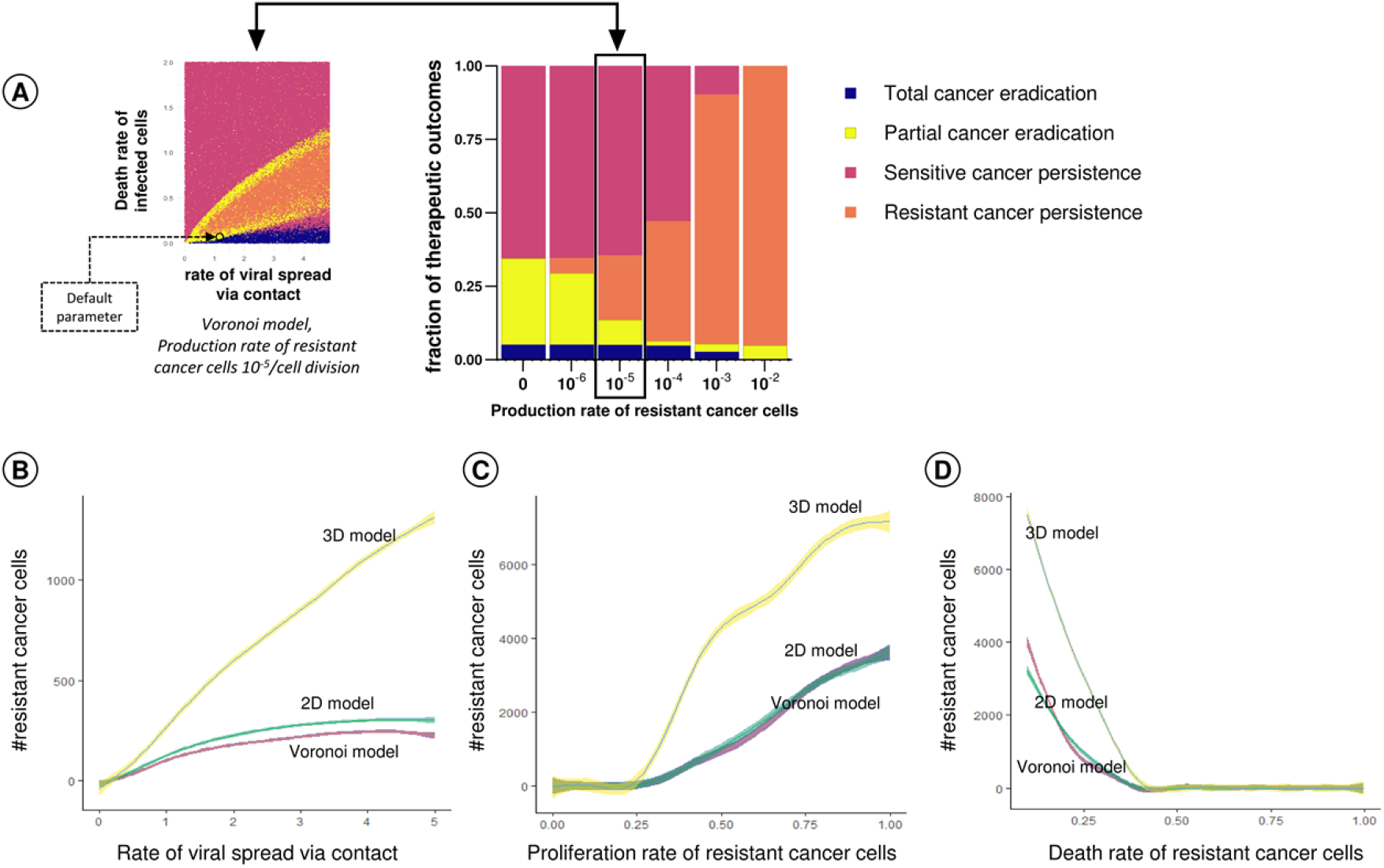
Factors influencing the persistence of resistant cancer cells in the tumor. (A) Effect of the production rate of virus-resistant cancer cells (*C*_r_) on the therapeutic outcome in the Voronoi model. For the same range of parameter values as in Fig 3 (rate of viral spread and death rate of infected cells) 100,000 simulations were run and classified as to their therapeutic outcome. The bar chart indicates the likelihood of the four outcomes for six values of *C*_r_. For ease of interpretation, the production rate of 10^−5^ per cell division is highlighted and linked to the Voronoi variant of Fig 3B. The size of the four bars in the bar chart is proportional to the areas indicated by blue, yellow, red, and orange in the panel to the left. Effects of (B) the rate *b*_i_ of viral spread, (C) the proliferation rate *b*_r_ of infection-resistant cancer cells, and (D) the death rate *d*_r_ of resistant cancer cells on the number of resistant cancer cells at the end of a simulation run. For graphs B to D, 10,000 simulations were run (for 1,000 parameter configurations and 10 replicates per configuration) for each of the three spatial configurations (2D grid, Voronoi, 3D grid), varying the parameter under investigation and keeping all other parameters at their default values (indicated by a black circle in figure 4A). The colored lines indicate the mean per parameter value, and the colored envelopes around the lines correspond to the 95% confidence band.

Figures 4BCD show how the “typical” number of virus-resistant cells at the end of the simulation depend on (B) the rate *b*_i_ of viral spread, (C) the proliferation rate *b*_r_ of infection-resistant cancer cells, and (D) the death rate *d*_r_ of resistant cancer cells (all other parameters were kept at their default values, as listed in Table 1 and indicated by a black circle in Fig.4A). The panels show that the conditions for the persistence of resistant cancer cells (*b*_r_ > 0.25; *d*_r_ < 0.45) are the same for the three spatial variants of the model (regular 2D, Voronoi 2D, regular 3D). However, the number of persisting resistant cells is much higher in the 3D model than in the two 2D models.

**Table 1:** Model parameters, their default values, and the range of values investigated in the simulations. Parameter values marked by a star are adapted from Berg et al. [15].

| Parameter | Explanation | Default value | Range |
| --- | --- | --- | --- |
| grid size | 10,000 cells ($100^2$ in 2D models, $22^3$ in 3D model) | 10,000 | 500 to 250,000 |
| $b_s$ | Proliferation rate of healthy stromal cells | 0.5* | - |
| $d_s$ | Death rate of healthy stromal cells | 0.2* | - |
| $b_c$ | Proliferation rate of infection-susceptible cancer cells | 1.0* | - |
| $d_c$ | Death rate of infection-susceptible cancer cells | 0.1* | - |
| $C_r$ | Production rate of resistant cancer cells per cancer cell division | $10^{-5}$ | $10^{-6}$ to $10^{-2}$, and 0 |
| $b_r$ | Proliferation rate of infection-resistant cancer cells | 0.9 | 0 to 1 |
| $d_r$ | Death rate of infection-resistant cancer cells | 0.2 | 0 to 1 |
| $F_i$ | Fraction of cells initially infected | 0.1* | - |
| $b_i$ | Rate of viral spread via contact of infected with susceptible cells | 1.2* | 0 to 25 |
| $d_i$ | Death rate of infected cells | 0.1* | 0 to 5 |
| $S_s$ | Relative susceptibility of stromal cells to viral infection | 0* | 0 or 1 |
| $P_{id}$ | Probability of a cell to be infected at a distance by an infected cell | 0* | 0 to 1 |
| $D_i$ | Distance of virus dispersal from an infected cell | 0* | 0 to 3 |
| $T_i$ | Time of virus introduction (infection) | 50 | 0 to 400 |
| $T_{max}$ | Total runtime of the simulation | 1,000* | - |

### Tumor progression and time of therapeutic intervention

Considering that tumor progression in time leads to an advanced stage of cancer and often accumulation of resistant cancer cells in the tissue, we assessed how the timing of virotherapy affects the therapeutic outcome. Figure 5A shows how the outcome depends on the time of virus introduction in the tumor (T_i_) in the 2D Voronoi model (see supplementary figure 7 for the regular 2D and the 3D model). In fact, there is not *one* outcome per introduction time T_i_ of the virus: replicate simulations for a given introduction time differ in therapeutic outcome, and Figure 5A represents the frequency distribution of these outcomes (for different introduction times, all other parameters were kept at their default values). When stromal cells are resistant to infection (top panel), the distribution of outcomes differs considerably depending on whether virotherapy starts before or after day 250. When the virus is introduced before day 250, total cancer eradication is only rarely achieved. Still, it is advantageous to start therapy as early as possible, as this maximizes the probability of partial cancer eradication. Interestingly, total cancer eradication is achieved in about 10% of the cases when virotherapy is started after 250 days. This can be explained by the fact that most stromal cells have disappeared when virotherapy starts at a late stage (figure 5B). As shown above, infection-resistant stromal cells may act as a spatial barrier, preventing the efficient spread of the virus in the tumor. It can therefore be beneficial to start virotherapy at a time when this barrier has disappeared (due to the fact that the stromal cells were outcompeted by the cancer cells). However, a late start of virotherapy has the downside that the persistence of a partially resistant cancer is the most likely outcome (figure 5A) and the number of resistant cancer cells is maximized (figure 5C). These negative effects of a late-starting virotherapy are more pronounced in the 3D variant of the model than in the two 2D variants (figure 5C, supplementary figure 7). Importantly, the effect of the time of treatment is stronger for other parameter combinations, most notably those resulting in partial cancer eradication and those resulting in the persistence of sensitive cancer due to resistance mediated by stromal cells (supplementary figure 8).

**Figure 5:**
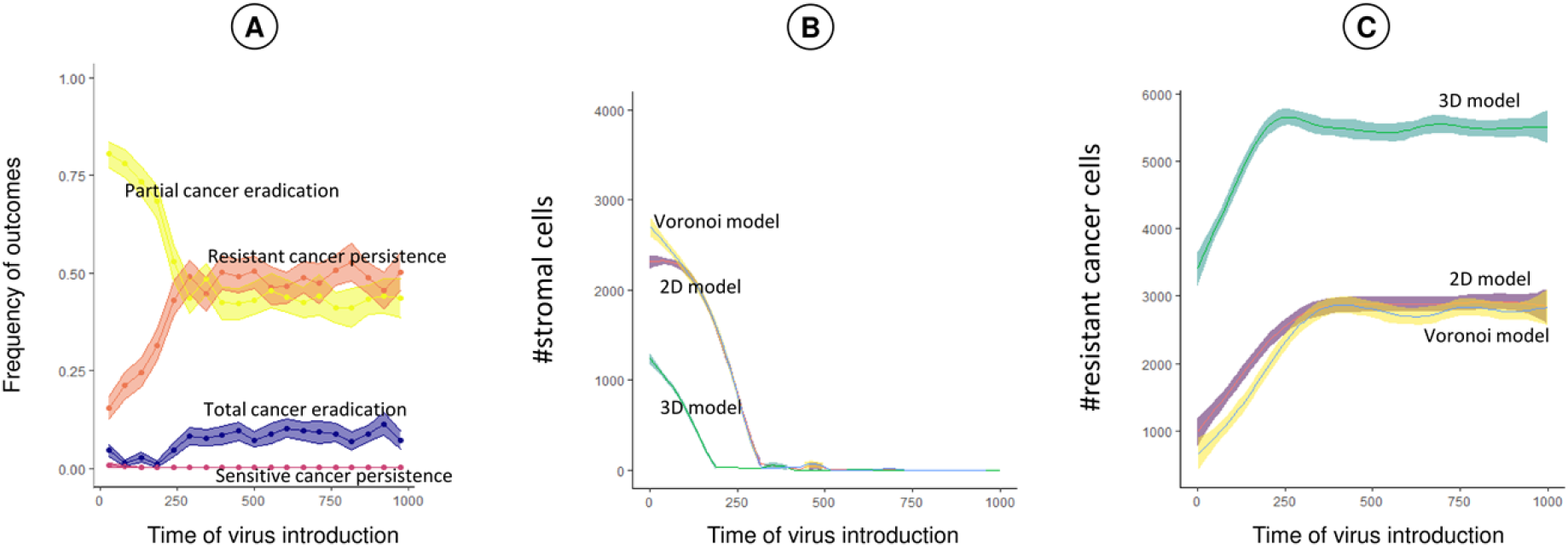
Effect of time of treatment on therapeutic outcome. (A) Effect of start of virotherapy (*T*_i_) on the therapeutic outcome in the Voronoi model and number of stromal cells (B) and number of infection-resistant cancer cells (C) at the end of the simulation. Each panel represents 10,000 simulations per spatial configuration: for 1000 virus introduction times *T*_i_ 10 replicate simulations were run, keeping all other model parameters at their default values. Due to stochasticity, the replicates for a given introduction time could differ in therapeutic outcome at time 1000. Colored lines indicate the mean value, colored envelopes indicate the 95% confidence band. Mean and confidence intervals in (A) were obtained via nonparametric bootstrapping.

### Dispersal of virus in the tumor

Up to now, we only considered contact-based transmission of the virus. If viral transmission only occurs between neighboring cells, it can be undermined by a barrier of virus-resistant cells (e.g. stromal cells). Figure 6 shows how the therapeutic outcome is affected if the virus cannot only spread via cell-to-cell contact but also via diffusion over small distances. The four panels in figure 6B show how, for our default parameters (indicated by a black circle in figure 4A) and the dispersal distances 0, 1, 2 and 3 (illustrated in figure 6A), the likelihood of the various therapeutic outcomes changes with an increase in *P*_id_, the probability that upon the death of an infected cell a given cell in the diffusion neighborhood of the dying cell is infected. The left-most panel in figure 6B (dispersal distance zero) considers the case where infection via dispersal does not occur and all virus transmission happens via cell-to-cell contact. As we have seen before, there are two therapeutic outcomes: partial cancer eradication (in about 75% of the simulations) and the persistence of a partially virus-resistant cancer (about 25%). Obviously, the outcome does not depend on *P*_id_ (as the diffusion neighborhood is empty). The three other panels in figure 6B illustrate how strongly the therapeutic outcome can be improved if viral diffusion is a potential form of viral transmission: already a dispersal distance of 2 or 3, virtually all simulations resulted in total cancer eradication; and even for a dispersal distance of 1, total cancer eradication was the most likely outcome for large values of *P*_id_.

**Figure 6:**
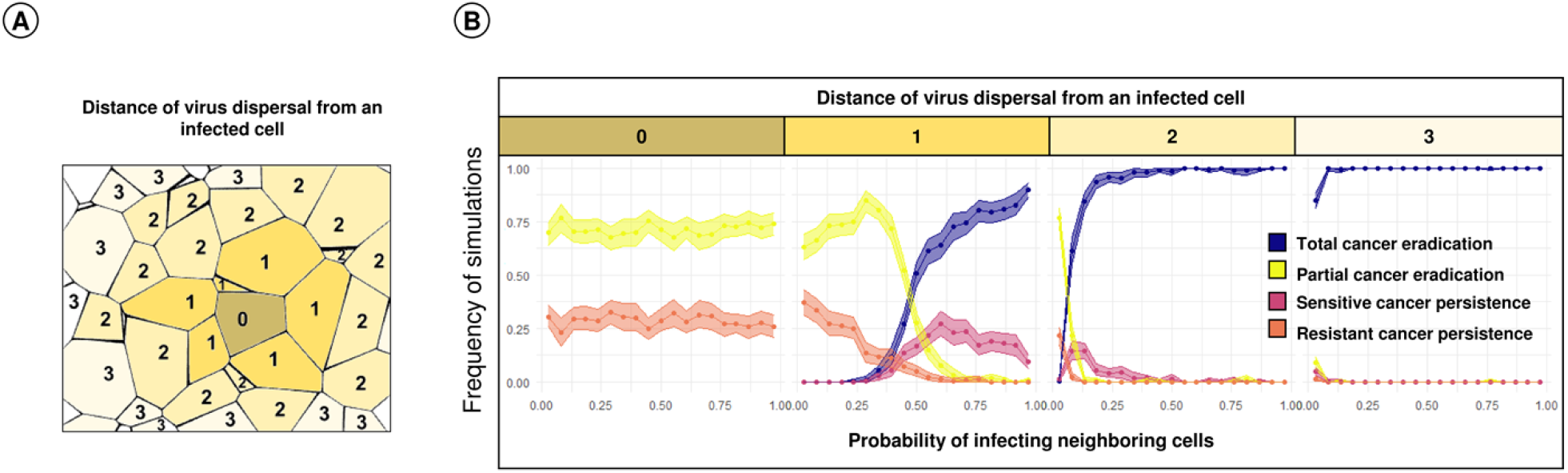
Effect of viral dispersal distance and the probability that neighboring cells are infected via diffusing viruses on the therapeutic outcome. In the Voronoi model, we considered various dispersal distances of the virus. For each dispersal distance, we changed the probability *P*_id_ that upon the death of an infected cell the cells in the diffusion neighborhood of that cell are infected, keeping all other parameters at their default values. (A) Interpretation of the viral dispersal distance, here ranging from 0 to 3. A focal cell marked “0” has distance zero to itself, distance one to its neighbors, distance two to its neighbors, etc. (B) Therapeutic outcome for dispersal distances 0, 1, 2, and 3 in relation to the probability *P*_id_ of being infected by an infected neighbor. For each graph, 10,000 simulations were run; colored lines and envelopes indicate the bootstrapped mean simulation outcome and the 95% confidence band.

## Discussion

The processes underlying therapeutic resistance to oncolytic viruses are so intricate that intuition and verbal reasoning can provide an incomplete and misleading account. Through a modelling approach, our study provides insight into the dynamics of oncolytic virotherapy and resistance within a spatial framework. Our results demonstrate that the outcome of virotherapy depends not only on the parameters governing virus replication and the spatial architecture of the tumor but also on the presence of resistant-cancer and -stromal cells that act as a barrier to the spread of oncolytic virus. More importantly, our results also provide an idea on how therapeutic outcomes can be improved by enhancing virus dispersal in the tumor and by sensitizing stromal cells towards virus infection. Overall, our model provides a systematic understanding of underlying parameters and their relative effect on the therapeutic outcomes of oncolytic virotherapy. For ease of access we provide executables of the model (see *data availability statement*), allowing users to challenge their insight and intuition by conducting a spectrum of *in silico* experiments themselves.

Our model is mainly intended as a conceptual tool that elucidates the intricate interplay of processes relevant for oncolytic virotherapy. Accordingly, we kept the model assumptions as simple as possible. Yet, we think that our results and conclusions are not unrealistic, as our model set-up and choice of parameters follows the experimentally validated study of Berg et al. [15]. It is reassuring that our model outcomes also align well with the experimentally validated study of Wodarz et al. [14], despite considerable differences in model structure (see supplementary figure 3 for details). In agreement with the studies of Berg et al. and Wodarz et al., we found that, for a given set of parameters, the therapeutic outcome is not deterministic but stochastic. For example, when we study the influence of rate of viral spread and death rate of infected cells (figure 3), we find that different outcomes are possible for neighboring sets of parameters. This may partly explain the variation in clinical outcomes. Stochastic effects are also evident in figure 5, where for the same time of virus introduction, various outcomes are possible, ranging from complete tumor eradication to persistence of resistant cancer.

This is the first theoretical study that explicitly addresses the implications of resistance against viral infection for oncolytic viral therapy. Our results suggest that even when resistance arises at a low frequency, such as at a rate of 10^−5^ per cell division, resistant cancer cells limit viral spread and prevent tumor eradication (figure 4A and supplementary figure 6). We also looked at rates ranging from 10^−6^ to 10^−2^, as not much information is available about the true occurrence of infection-resistant cells in the tumor. However, this range seems reasonable as we find at least 1-10% infection-resistant cells in patient-derived cancer cell-lines [8,23]. We assume that resistance to viral infection may come at a metabolic or physiological cost that reduces the rate of proliferation or increases the death rate of resistant cancer cells. This is justified by the experimental observations that production of antiviral factors and related signaling pathways can cause arrest of cell-cycle thereby decreasing proliferation rate [24] and may also lead to stress-induced cell death [25,26]. Not surprisingly, our simulations indicate that a high cost of resistance is associated with a poor survival of resistant cancer cells and leads to a significant decrease in their frequency (figure 4C-D). The cost of resistance has a significant influence only in scenarios where the tumor persists due to selection of resistant cancer cells. “Perhaps more surprisingly”, our results indicate that in the scenario with a faster rate of viral spread there is a rapid clearance of infection-sensitive cancer cells, which in turn promotes the selection and survival of resistant cancer cells. Indeed, this has also been found to be true in practice where therapeutic intervention causes selection of resistant clones [27].

Our study highlights the importance of stromal cells for oncolytic virotherapy. Even if stromal cells are not affected by the presence of the virus (as we assume in most of our simulations), they can prevent viral spread as well. Hereby they can contribute to therapy failure when viral infection and viral replication is specific to cancer cells only. Our observations from the simulations are in line with various experimental studies that observe an improvement in therapeutic outcomes upon sensitizing normal cells or upon using viruses that are not only specific for cancer cells [28] [29,30]. We observe that sensitizing stromal cells to viral infection increases the chances of the virus to spread from an infected cell to non-infected (sensitive) cancer cells in the neighborhood, which results in an improvement in the efficacy of tumor eradication. However, sensitizing stromal cells does not affect persistence of resistant cancer cells and comes at a cost of stromal cell death and possible cytotoxicity to the healthy tissue present in the microenvironment (supplementary fig5). Resistant stromal and cancer cells thus prevent cell-to-cell transmission of virus in the tumor where we find that failure during early therapy occurs due to presence of resistant stromal cells and during late therapy occurs due to presence of resistant cancer cells (figure 5).

However, not all oncolytic viruses depend on a cell-to-cell mode of transmission and instead, can infect cells by diffusing through extracellular space. Modeling such a diffuse mode of infection, we find that extending the diffusion of virus particles even by a single degree of neighbor can significantly increase the number of outcomes that result in partial or complete tumor eradication. Our findings are in agreement with the observations that an improved diffusion coefficient increases the number of infected cells in the vicinity [31], and that the probability of infection can depend on neutralization by antibodies and frequency of infectious virus particles released upon cell lysis [6,32]. Again, these results highlight the importance of explicitly considering spatial features and extracellular factors in order to gain insight in understanding virus-tumor interactions.

Our model addressed the importance of spatial configuration and size of the simulation grid (total cell number). We find that the grid size has an impact on the therapeutic outcomes; where simulations running in smaller grids (500 cells) lead to partial cancer eradication (indicated by a wider ‘yellow’ area in supplementary figure 1) instead of persistence of a resistant cancer (‘orange’ area). Due to a higher number of total cells in a larger grid, the probability of generation of resistant cancer cells is higher even if they arise at a low frequency (10^5^) per cell division. Nevertheless, the overall results are robust and comparable between grids of different sizes that contain at least 10,000 cells in total. Considering the spatial configuration of the simulation grid, we find that in 2D, both the Voronoi (three to six neighbors) and the regular model (four neighbors) yielded very similar results, despite the difference in connectivity between cells. This is in agreement with the conclusions of Berg et al. [15], who also found strong similarities between regular and Voronoi grids. However, when including a third dimension, we find that (sensitive or resistant) cancer cells were able to persist in a much larger parameter space. Such differences between 2D and 3D settings could hint at potential discrepancies when comparing *in vitro* results obtained in a (more or less) two dimensional petri-dish environment, and *in vivo* results obtained in a three dimensional setting [15,33,34]. One might assume that the differences in outcomes could result due to a significant difference in the number of boundary cells in the 2D model (5% of the total cells) or 3D model (25% of the total cells) in a grid of 10,000 cells. However, this is unlikely because we do not find any differences in outcomes upon changing the grid-size of the 2D or 3D models (supplementary figure 1) as grid-size could directly influence the number of boundary cells. Alternatively, it might be possible that the outcomes differ between the spatial configurations due to the differences in the number of neighbor cells. However, we do not find any differences between the outcomes resulting from the two 2D models even though there is a difference in the number of neighboring cells (4 in case of regular and 4-6 in the case of Voronoi). On the other hand, the Voronoi and the 3D model have a similar number of neighboring cells but do differ in the outcomes. This suggests that in the 3D model, as cells occupy different planes in the grid, there is an extra degree of freedom for the cancer cells to escape or even for the virus to spread. This may increase the chances of partial tumor eradication, which is something that we observe in our model (supplementary figure 1).

To minimize the complexity of our model, we have neglected intra-tumoral heterogeneity. However, this can be easily incorporated in the model by introducing different cancer cell-clones that vary in their rates of proliferation and death and sensitivity to viral infection. Additionally, one may also implement a degree of sensitivity to viral infection and resulting changes in the rate of cell-death to have a finer understanding of the virus-cancer dynamics. Most importantly, we have neglected the effect of immune response during virotherapy. This will be the subject of a future study where we aim to model the anti-tumor and anti-viral immune responses in a similar spatial configuration.

Current experimental research has demonstrated the possibility of employing effective oncolytic virotherapy candidates in clinics; however, therapeutic resistance will remain to be a challenge unless efforts are made to understand the underlying mechanisms. We hope that our computational approach aids in defining the impact of the various factors that may influence resistance and thereby therapeutic efficacy of virotherapy. We are confident that our results, along with experimental observations, can assist the scientific community in improving the design of virotherapy.

## Acknowledgements

We thank Prof. Dr. Roger Chammas from the University of São Paulo for useful discussions and the Center for Information Technology of the University of Groningen for their support and for providing access to the Peregrine High Performance Computing cluster.

## Data Availability Statement

To improve usability, we provide two different versions of the model for all common operating systems: 1) a terminal only version, which reads parameters from a configuration file and which can be used to perform demanding simulations on a high performance computation cluster and 2) a graphical version, where the user is provided with an intuitive graphical user interface and can visually observe the spatial interplay between cell types. The code used for this work and an executable version of the Oncolytic Virus Resistance simulator (OVR) can be found in the Supplementary data or on www.github.com/rugtres/ovr.

## Funding

The work of FJW is supported by the European Research council (ERC Advanced Grant No. 789240).

D.K.B. received a PhD scholarship from CAPES (Finance Code 001) and ATTP-GSMS (Abel Tasman Talent Program) Groningen.

## Competing Interests

The authors declare that there are no competing interests.

## Author contributions

Conceptualization: DKB, TJ, TD, FJW; Implementation: TJ, DKB; Model analysis: DKB; Writing: DKB, TJ, TD, FJW

## Supplementary Figures

**Supplementary figure 1:**
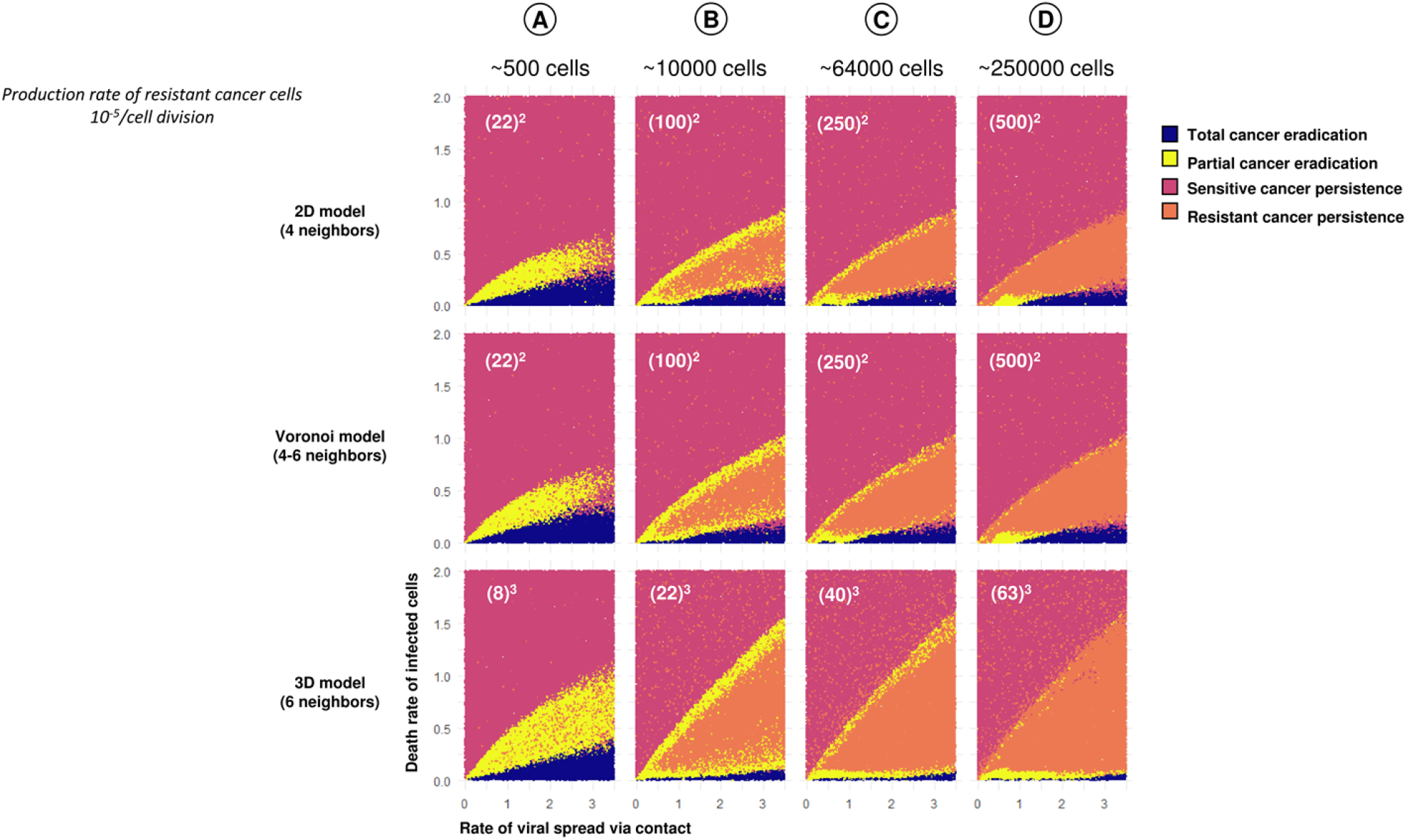
Effect of grid size and spatial configuration on the outcome of virotherapy. Therapeutic outcomes in relation to the rates of viral spread via contact and death rate of infected cells for the three spatial configurations considered (regular 2D grid, 2D Voronoi model, regular 3D grid) and for various population sizes (A) about 500 cells; (B) about 10,000 cells; (C) about 64,000 cells; (D) about 250,000 cells. The white text in each panel indicates how the population size relates to the grid dimensions. 50,000 simulations were run for each panel, and each point corresponds to one simulation. With the exception of population size, all parameters were at their default values.

**Supplementary figure 2:**
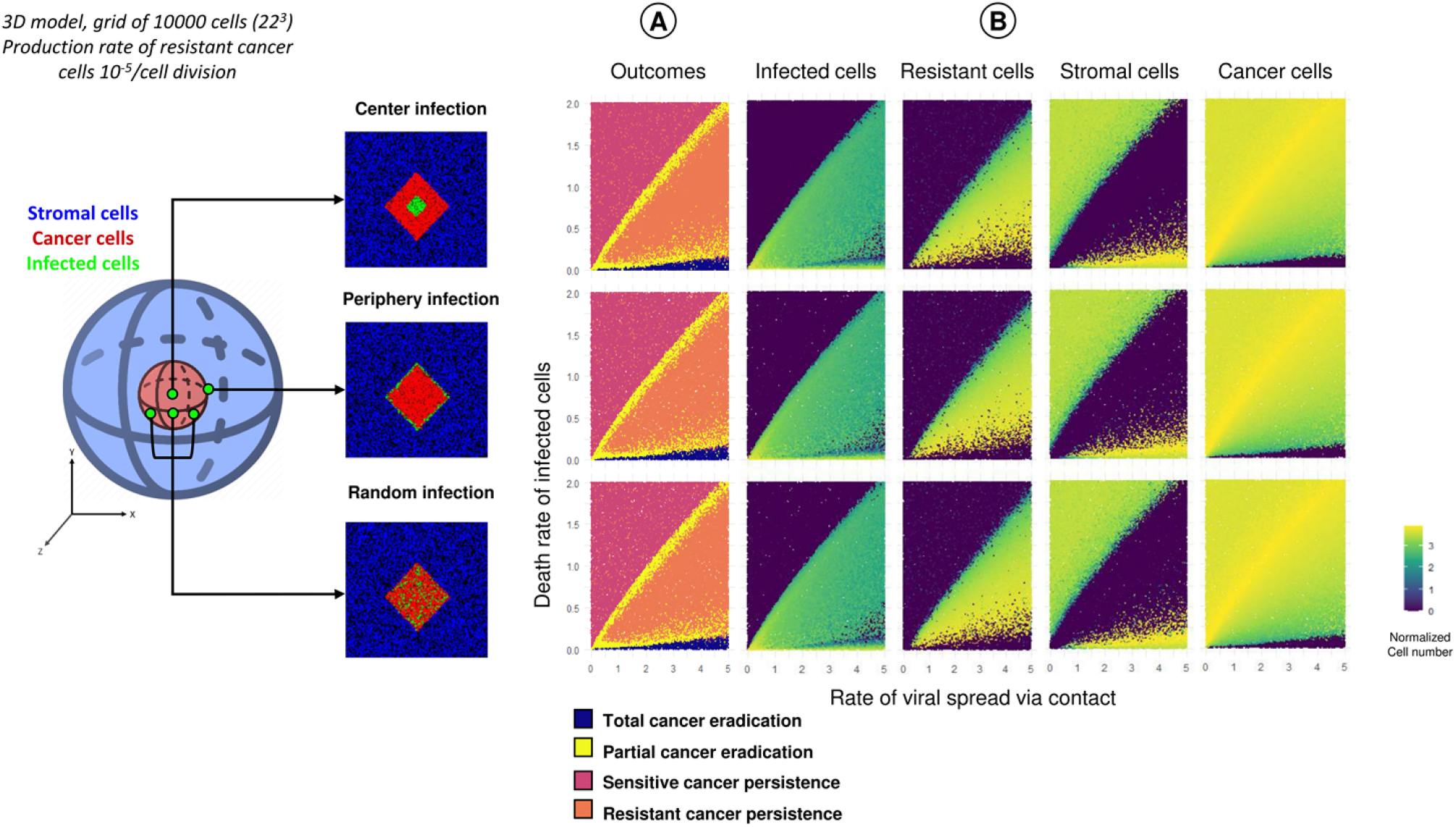
Effect of different forms of viral infection on the therapeutic outcome. Simulation outcome in relation to the rates of viral spread via contact and death rate of infected cells in the 3D model for three different forms of viral infection. As indicated in the illustration on the left, virus infection in tumor is initiated either in the center (top row), or from the periphery (middle row), or in a random manner (bottom row). For each infection scenario 50,000 simulations were run, which were classified according to (A) their therapeutic outcomes; and (B) the number of the different types of cells at the end of the simulations. All parameters were at their default values. The color code is based on the logarithm of cell numbers.

**Supplementary figure 3:**
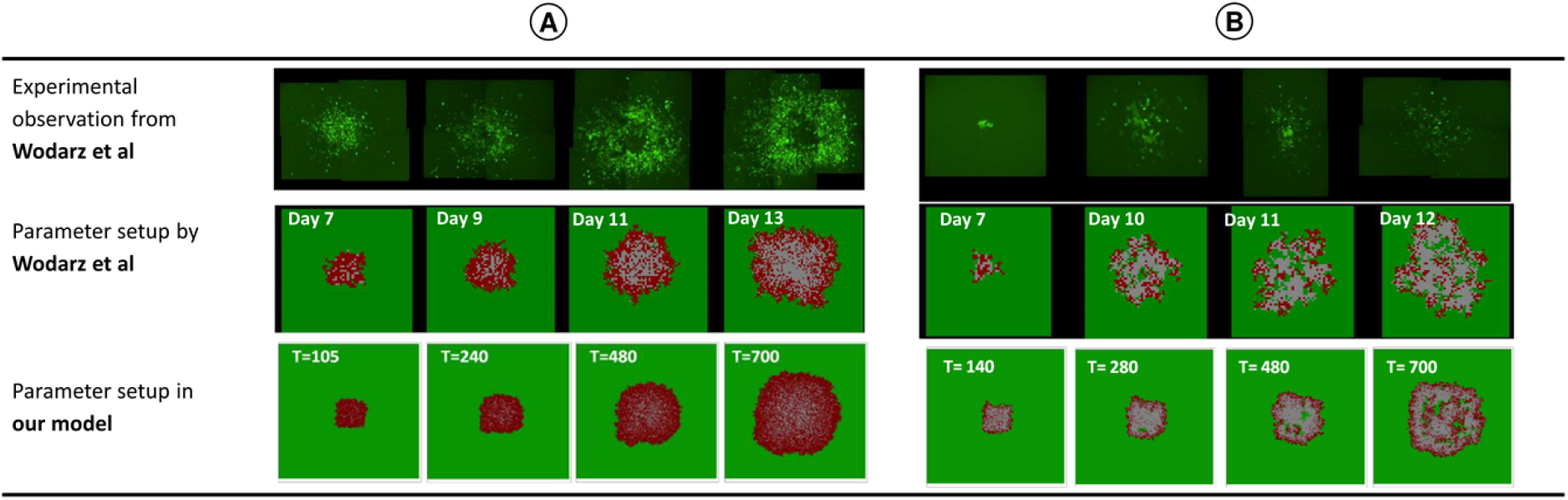
Comparison of our model with the results of Wodarz et al. (2012). In their Figures 4 and 5, Wodarz and colleagues [14] compare the spatial dynamics of experimentally induced viral infections with the predictions of an agent-based model. Here, we illustrate that our event-based model creates similar spatial patterns as the discrete-time Wodarz model. (A) In both models, virotherapy leads to the radial spread of the virus and the extinction of cancer cells if the viral infection rate exceeds the death rate of infected cells by a factor of at least 10. In the simulation shown, *b*_i_ = 0.1 and *d*_i_ = 0.001. (B) In both models, virotherapy results in the long-term coexistence of infected and uninfected cancer cells if the viral infection rate does neither exceed the death rate of infected cells nor the birth rate of uninfected cells by a factor of 10. In the simulation shown, *b*_i_ = 0.1, *b*_c_ = 1.0 and *d*_i_= 0.01. In the experimental observations of Wodarz and colleagues, infected cells are labelled as green and non-infected cells as black. Regarding the simulations, we follow the conventions of Wodarz and colleagues to label infected cancer cells as red, non-infected cells as green, and empty space as white.

**Supplementary figure 4:**
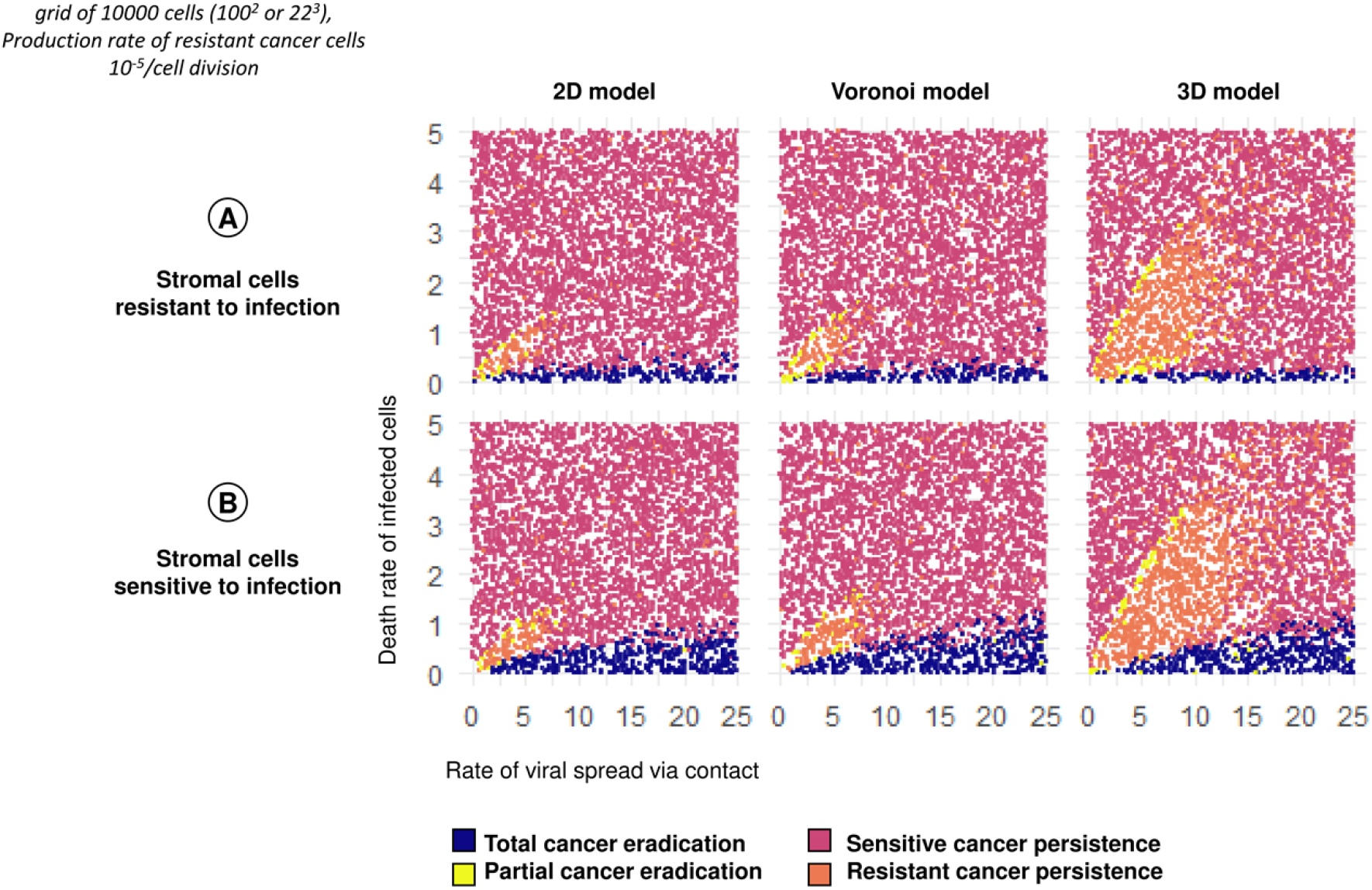
Therapeutic outcome for an increased parameter range of rate of viral spread via contact and death rate of infected cells. In the graphs of the main text (figure 3), the rate of virus spread ranges from 0 to 5, while the range of death rate of infected cells from 0 to 2. For the three spatial configurations considered (regular 2D grid, 2D Voronoi model, regular 3D grid), the panels show the therapeutic outcome for a wider range of parameters, for two scenarios: (A) stromal cells cannot be infected by the virus; (B) stromal cells are sensitive to infection. Each panel in the figure is based on at least 10,000 simulations and each point represents one simulation. All parameters that are not varied are at their default values.

**Supplementary figure 5:**
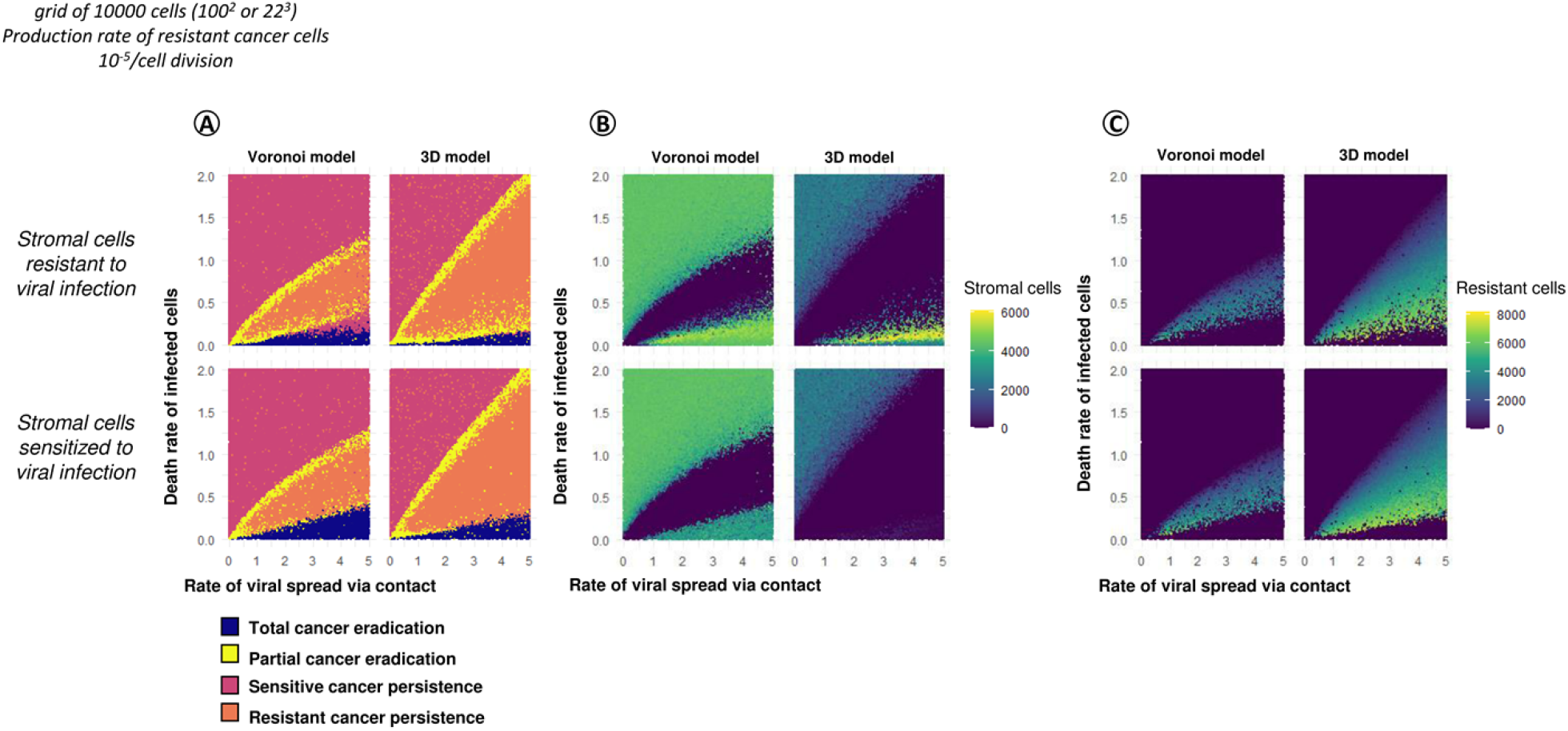
Consequences of sensitizing stromal cells for virotherapy. (A) Therapeutic outcome in relation to the rates of viral spread via contact and death rate of infected cells for two spatial configurations (2D Voronoi model, regular 3D grid) and two different assumptions on stromal cells: stromal cells cannot be infected by the virus (top row); or stromal cells are sensitive to infection (bottom row). Each of the four panels represents 100,000 simulations. For the simulations in (A), the four panels indicate the number of stromal cells (B) and the number of resistant cancer cells (C) at the end of the simulation. The color code is based on the absolute of cell numbers.

**Supplementary figure 6:**
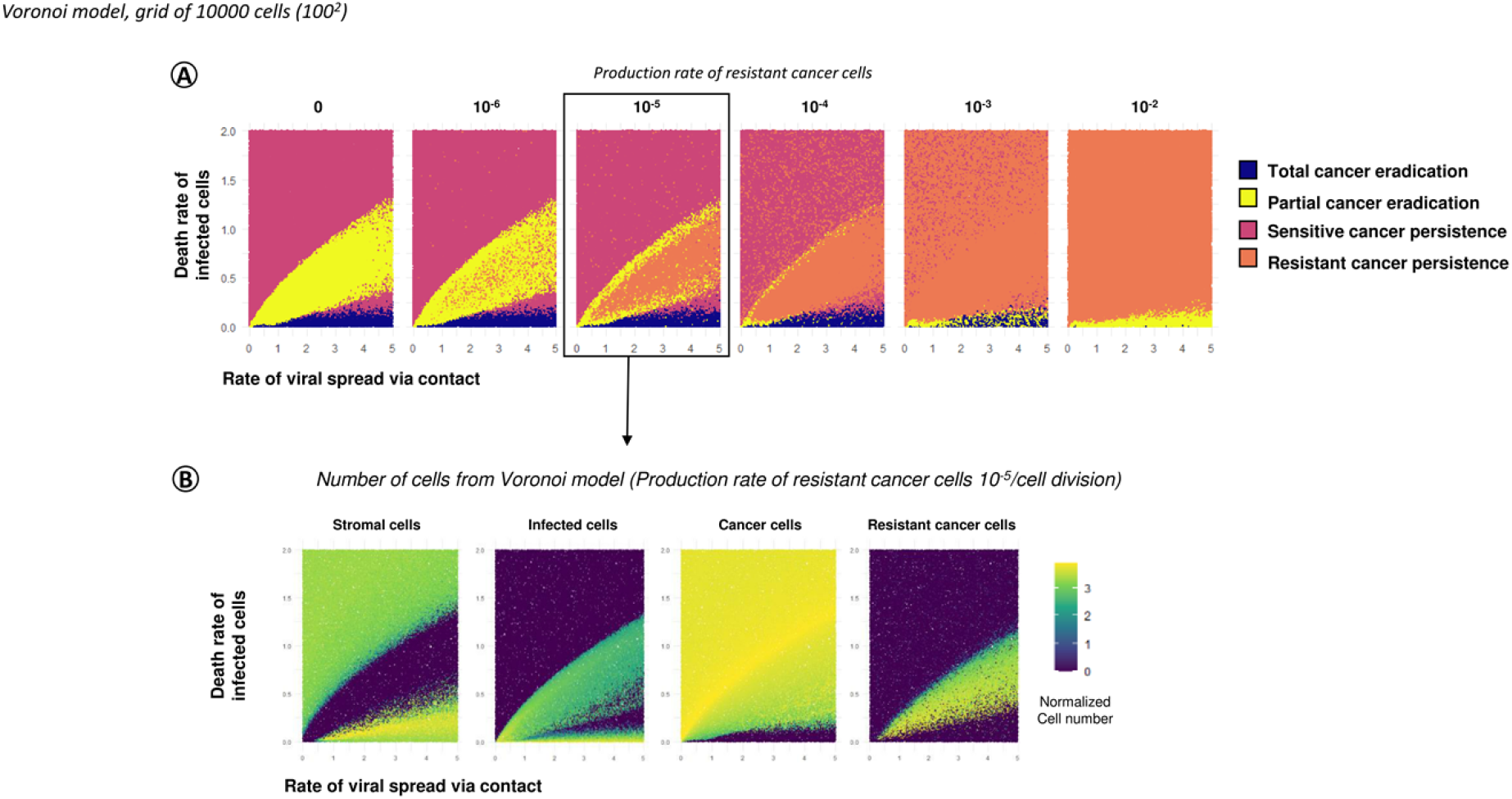
Effect of the production rate of virus-resistant cancer cells on the simulation outcome. (A) Therapeutic outcomes in the 2D Voronoi model in relation to the rates of viral spread via contact and death rate of infected cells for six production rates of virus-resistant cells (ranging from 0 to 10^−2^ per cell division). Each panel represents 100,000 simulations, and each point corresponds to one simulation. (B) Numbers of different types of cells at the end of the simulation for the default production rate of 10^−5^ per cell division. The color code is based on the logarithm of cell numbers.

**Supplementary figure 7:**
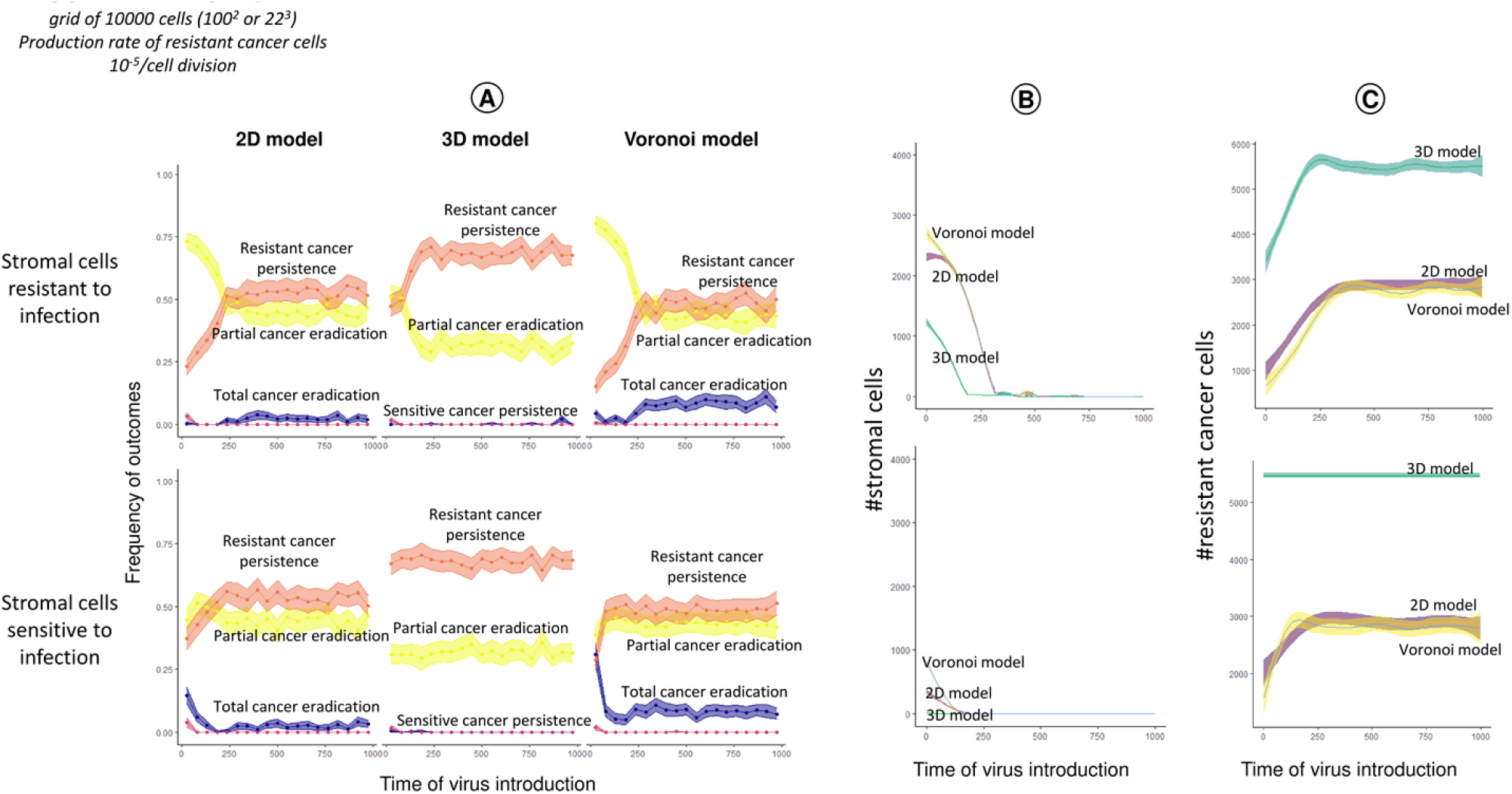
Effect of time of treatment on therapeutic outcome. This figure corresponds to Fig 5 in the main text, which illustrates the effect of start of viral treatment on the therapeutic outcome in the 2D Voronoi model. Here, the corresponding outcomes are shown in (A) for the regular 2D grid and 3D grid models. Effect of start of virotherapy on the (B) number of stromal cells and (C) number of infection-resistant cancer cells at the end of the simulation is provided. Two scenarios are considered: stromal cells are resistant (top row) or sensitive (bottom row) to infection. The number of simulations per panel and the model parameters are as in Fig 5.

**Supplementary figure 8:**
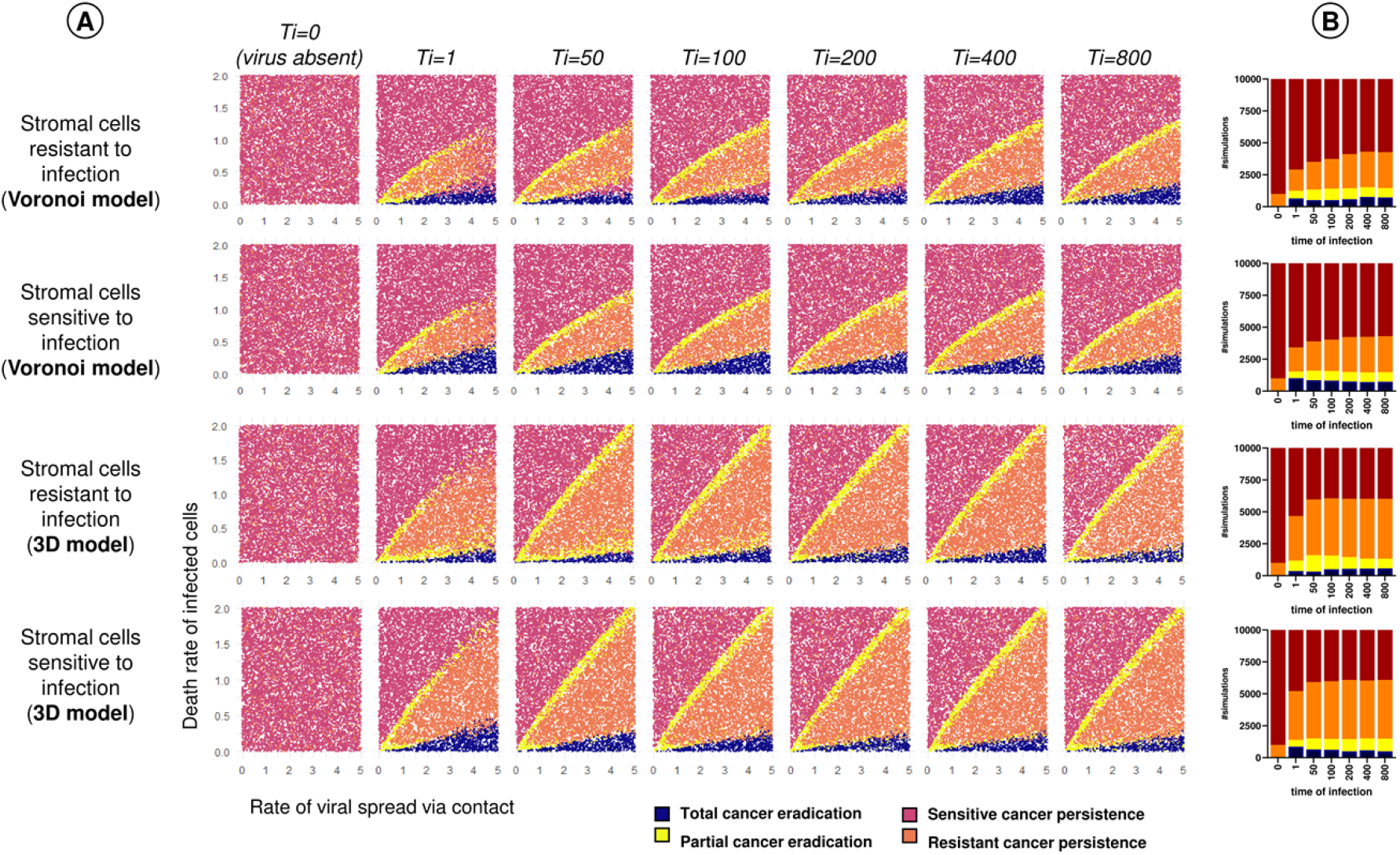
Effect of time of treatment with respect to rate of viral spread and death rate of infected cells on therapeutic outcome. (A) Effect of the time of treatment (*T*_i_) on the therapeutic outcome in the Voronoi model and 3D model. For the same range of parameter values as in Fig 3 (rate of viral spread and death rate of infected cells) 10,000 simulations were run and classified as to their therapeutic outcome. (B) The bar chart indicates the likelihood of the four outcomes for different values of *T*_i_.

## Notes

### Competing Interest Statement

The authors have declared no competing interest.

